# Molecular heterogeneity and differential metabolic signatures in thymic adipocytes

**DOI:** 10.64898/2026.01.14.699556

**Authors:** Joon Schwakopf, Amber R. Syage, Anca Franzini, Katherine E. Varley, Dean Tantin

## Abstract

Adipocyte depots throughout the body are physiologically and molecularly distinct. With age, adipocytes increase in and around aged thymi. However, thymic adipocytes completely lack molecular characterization. We developed and optimized methods to isolate adipocyte nuclei from mouse thymi of different ages and sexes. Single-nucleus multiomic analysis of male and female mice aged 4-9 months reveals that thymic adipocytes are heterogeneous, with at least two distinct populations. One subpopulation harbors a transcription and chromatin signature consistent with beige/brown fat. A larger subpopulation more strongly resembles classic white adipose tissue and expresses genes associated with epithelial-to-mesenchymal transition (EMT) and antigen presentation. Analysis of differentially open chromatin in the white compared to beige adipose population identifies binding sites for Foxn1 and HIF-1α/Arnt, consistent with a situation in which thymic white adipose cells emerge from thymic epithelial cells, possibly under hypoxic conditions. Immunofluorescence microscopy confirmed the expression of UCP1 protein in cells within the thymic parenchyma, most prominently in subcapsular cortical regions. This resource reveals a complex milieu of thymic adipocytes and identifies multiple avenues for directly probing their ontogeny, dynamics and functional significance.

## INTRODUCTION

The thymus is the primary site of T cell development and education and thus plays a central role in adaptive immunity and central tolerance ^1^. It is a complex organ that consists of developing T cells (thymocytes), thymic epithelial cells (TECs), fibroblastic stromal cells, macrophages, dendritic cells (DCs), B cells (particularly in the medullar), mature nonconventional T cells (e.g., iNKTs), endothelial cells, pericytes, nerves, and other less abundant cell types. The aging thymus also accumulates abundant and dynamic population of adipocytes ^1^, which we here term ThyAds.

With age, areas of the thymus devoted to thymopoiesis atrophy and T cell output declines in a process termed thymic involution ^2,3^. Involution is associated with progressive disruption of the cortico-medullary junction, loss of thymic output, and increased adiposity in a manner influenced by biological sex and diet ^4^. Attenuation of thymic output contributes to reduced TCR diversity, impaired pathogen response and increased susceptibility to infection ^1,5–9^. Additionally, involution can be accelerated by administration of thiazolidinediones, implicating adipocyte expansion in this process ^10^, however their function in this process is unclear. Thymic involution is conserved across vertebrates and recapitulated in laboratory mice. Although severe infection, radiation, malnutrition, or hormonal alterations can induce acute thymic atrophy, age-related involution is the setting in which adipocyte accumulation is most consistently observed ^1,4,11–13^. Despite their increasing abundance in the aging thymus, and the possibility that they have specialized functions analogous to other tissue-specific fat depots ^14^, ThyAds have yet been directly studied at the molecular level. Prior studies focused on thymocytes and thymic stromal cells have omitted ThyAds because intact adipocytes are large, buoyant, and fragile, and are therefore poorly captured by droplet-based microfluidic workflow ^15–18^.

Recent single-nuclei methodologies combining RNA-seq and ATAC-seq have enabled high-resolution mapping of adipose depots, including age-associated pro-inflammatory and metabolic shifts linked to obesity and related pathologies. These technologies have also shown that adipose depots are tailored to their local environment and differ in cellular and molecular composition across the body ^14^. For example, subcutaneous and visceral WAT have distinct developmental origins, mechanisms of expansion and disease associations ^19^. Additionally, adipocytes remodel over time ^20,21^. In humans highly metabolically active brown adipose tissue (BAT), subcutaneous white adipose tissue (WAT), and beige fat, an intermediate form, typically decline with age whereas visceral WAT expands ^22–24^. Together, these findings led us to hypothesize that ThyAds are distinct from other adipocyte populations and that their age-associated expansion reflects changes in cellular composition and state.

Here, we used an integrated multiomic strategy to generate the first single-nuclei dataset of ThyAds. By profiling nuclei rather than intact cells, multiomic sequencing overcomes the technical limitations imposed by adipocyte size and fragility. We develop methodologies to efficiently liberate and isolate ThyAds and their nuclei from mouse thymi. By single-nucleus profiling, we identify two highly distinct ThyAd populations, termed ThyAd I and ThyAd II. ThyAd I exhibit a beige fat-like transcriptional program, whereas ThyAd II show features consistent with classic WAT. Notably, ThyAd II-specific accessible chromatin was enriched for consensus sites recognized by Foxn1, the master transcription regulator of TECs. This enrichment suggests a TEC-linked regulatory signature and supports the hypothesis that ThyAd II derive from TECs. Together, our ThyAd multiome resolves two adipocyte programs and provide clues to cellular origin, providing a roadmap to test lineage relationships and functional roles in thymic aging.

## RESULTS

### ThyAds comprise two discrete subpopulations, one of which has transcriptional hallmarks of beige fat

Thymic adipocytes are technically difficult to recover. We successfully developed an enrichment workflow that combines a retina tissue dissociation protocol and WAT buoyancy-based fractionation, followed by single-nuclei isolation to profile 6-month-old female mice (**Figure 1A**). In mature thymi, scattered multilocular lipid-laden cells are detectable within the subcapsular region, parenchyma, and PVS by ∼3 months, while large unilocular white adipocytes become prominent after ∼12 months ^25^. We therefore reasoned that 6-month-old mice would capture maximal diversity in the ThyAd pool, at a timepoint when thymic output is still preserved but ThyAds are recoverable from a sufficient amount of thymic tissue. After dissection, we extensively cleaned the thymic capsule to minimize contamination while preserving subcapsular and septal ThyAds (**Figure 1B**). Thymi were mechanically and enzymatically digested to generate a single-cell suspension, then fractionated to collect the buoyant adipocyte-enriched layer. Nuclei prepared from the buoyant fraction were robust, with preserved euchromatic and heterochromatic regions and nucleoli (**Figure S1A**).

**Figure 1.**
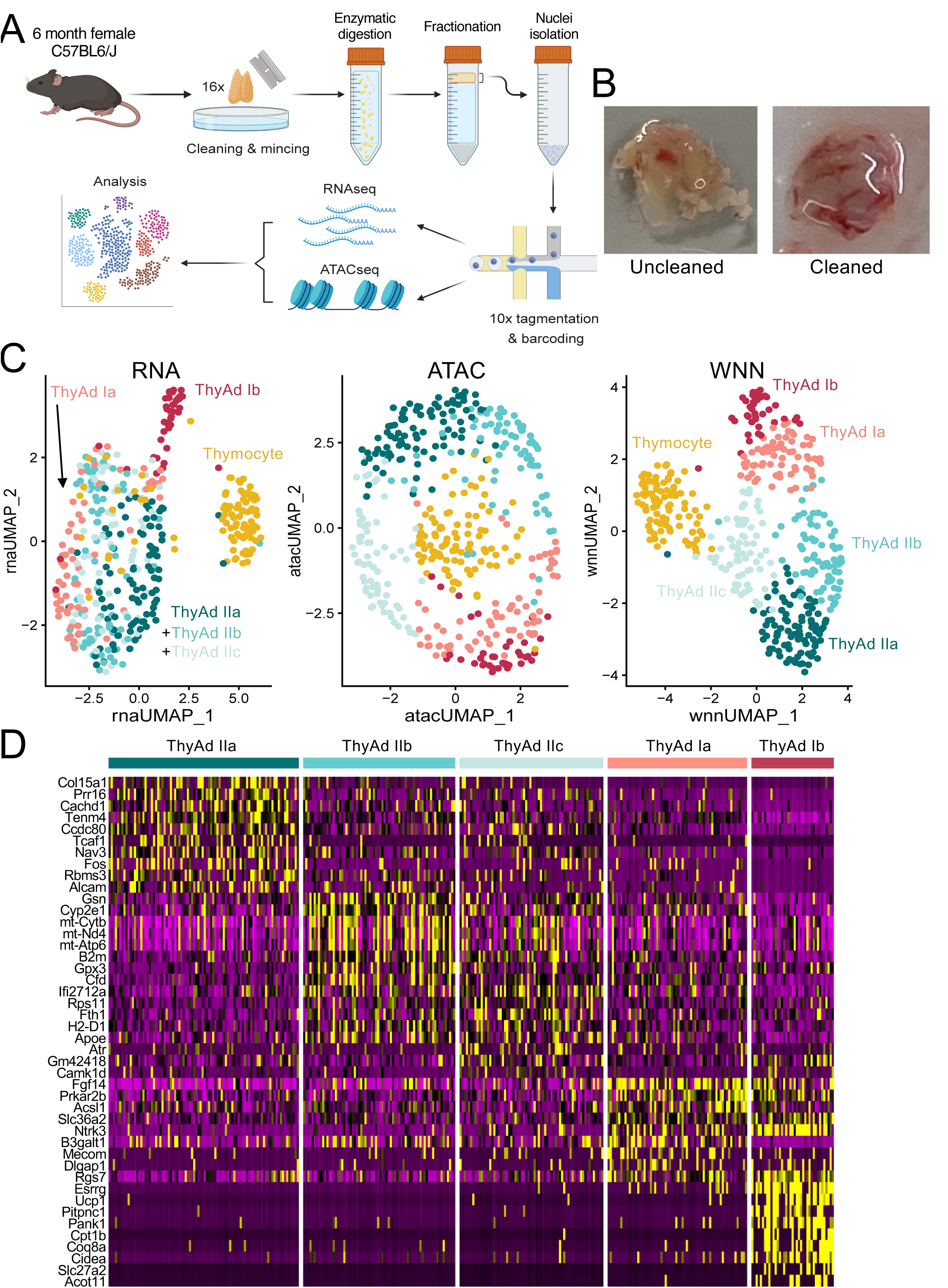
Female ThyAds are heterogeneous, with one subclustered harboring transcriptional signatures of metabolically active beige fat. (A) Experimental design for isolation and multiomic sequencing of murine ThyAds. Thymi were processed by mechanical disaggregation, digestion, centrifugation (isolation by buoyancy) and nuclei isolation prior to single-nucleus ATACseq + RNAseq multiomics. (B) Example thymus prior to and after removal of perithymic fat. The septa and thymic lobes were kept intact such that the fatty tissue between the lobes is retained. (C) UMAP projection and clustering of ThyAds by RNAseq, ATACseq, and Weighted Next Neighbor (WNN) analysis. ThyAds (n = 429, avg gene count 2,974) were reclustered after removal of Prprc-expressing nuclei. A residual Ptprc+ cluster was also present. (D) Expression of most distinguishing ThyAd marker genes by cluster. A heatmap displaying the top 10 DEGs within each ThyAd subpopulation is shown.

16 pooled female thymi were used for single-nucleus ATAC-seq plus RNA-seq, yielding 3,610 high-quality nuclei after filtering (**Figure S1B**). Initial uniform manifold approximation and projection (UMAP) clustering indicated substantial hematopoietic carryover, with most nuclei expressing *Ptprc* (*Cd45*) and resolving into expected thymocyte populations based on canonical markers such as *Cd8a*, *Cd4*, *Mki67*, *Hes1* and *Cd5* (**Figure S1C,D**). For example, cluster 8 expressed *Il2ra* but not *Cd8a* or *Cd4*, identifying these cells as DN2/3. Cluster 2 expressed *Cd5* and likely comprised B cells. Cluster 7 expressed *Klrb1c*, which encodes the NK1.1 surface protein. These nuclei are likely derived from iNKT cells. In contrast, signatures of non-hematopoietic thymic stromal populations (TECs, fibroblasts/mesenchyme, endothelium) were minimal or absent. This may reflect either or both a greater abundance of bone marrow-derived cells in the thymus, or a closer apposition of these cells with ThyAds such that they are preferentially carried over with the buoyant fraction following digestion. These data establish a technical framework for ThyAd recovery and set the stage for targeted analysis of the ThyAd compartment.

Two transcriptionally discrete clusters (3 and 10) expressed the canonical adipocyte markers *Pparg* and *Plin1* (**Figure S1C**) as well as *Fabp4* (not shown), consistent with adipocytes. A lipid metabolism gene module further supported adipocyte identity in clusters 3 and 10, whereas cluster 11 (the least abundant cluster) showed weaker and less consistent expression of adipocyte-associated transcripts (**Figure S2A**). We therefore focused on adipocyte clusters 3 and 10, yielding 429 high-quality adipocyte nuclei. These nuclei contained a mean of 1533 detected genes and an average depth of 2974 reads per nucleus.

We re-clustered adipocyte based on gene expression (RNA-seq UMAP), chromatin accessibility (ATAC-seq UMAP), and a weighted nearest neighbor (WNN) approach that integrates both modalities (**Figure 1C**). This analysis resolved two broad transcriptional programs, termed ThyAd I and ThyAd II, distinguished by differential expression of lipid metabolic genes (**Figure S2B**). A small residual thymocyte cluster expressed *Themis* and *Satb1* (**Figure S3A**) as well as lipid metabolic genes such as *Fasn* (**Figure S2B**) and may reflect doublet nuclei. The remaining clusters lacked thymocyte gene expression and robustly expressed adipocyte markers (e.g., *Pparg*), supporting their adipocyte identity (**Figures S2B** and **S3A**).

To define transcriptional states within ThyAds, we identified the top 10 DEGs across clusters. This revealed clear separation between ThyAd transcription programs, with ThyAd I more strongly expressing genes associated with thermogenesis relative to ThyAd II (**Figure S3A**). Restricting analysis to adipocyte nuclei only, we further resolved five distinct ThyAd subclusters–Ia, Ib, IIa, IIb and IIc (**Figure 1D**), consistent with prior evidence showing that ThyAds are morphologically heterogeneous ^25^.

ThyAd Ia nuclei were marked by the expression of Prkar2b and *Fgf14*. In contrast, ThyAd Ib more robustly expressed *Esrrg*, *Ucp1*, and *Cidea* (**Figure 1D**). *Ppara* was also up-regulated in ThyAd Ib (**Table S1**). These genes have been shown to positively regulate beige adipocytes ^26–28^. These findings strongly suggest that ThyAd I – particularly ThyAd Ib – are beige-like, consistent with work that used imaging to identify UCP1-positive cells detected inside the thymic parenchyma ^25^. In contrast, ThyAd II nuclei expressed a distinct program consistent with WAT-like identity (e.g., *Apoe*, **Table S1**) and additional features no observed in ThyAd I. ThyAd IIa was enriched for epithelial-mesenchymal transition (EMT)-associated transcripts including *Prr16* and *Alcam*, while ThyAd IIb/IIc displayed higher levels of *B2m* and *H2-D1* (**Figure 1D**), consistent with increased MHC-I-related gene expression ^29^. ThyAd II also expressed *Gpx3*, which has been reported in thymic stromal populations including intermediary fibroblast-like states ^30^. Additional feature plots are shown in **Figure S3B**. These results indicate that ThyAd I (in particular ThyAd Ib) resembles beige-like adipocytes, whereas ThyAd II is more WAT-like and exhibits an EMT-/antigen-presentation-associated transcriptional signature.

Because ThyAd I and ThyAd II subclusters were readily distinguishable by the thermogenic transcription signature in ThyAd I (**Figure 1D**), we combined the ThyAd Ia+Ib and IIa+IIb+IIc subclusters and performed differential expression analysis (**Table S2**). Gene expression differences between combined ThyAd I and ThyAd II nuclei are shown in **Figure 2A**. For example, *Prdm16* and *Ppara* were significantly enriched in ThyAd I, encoding transcriptional regulators of beige/brown fat ^31,32^, while Retn and Prr16 were significantly enriched in ThyAd II, encoding WAT-associated protein Prr16 ^33^ and the adipokine resistin, a regulator of adipogenesis ^34^, respectively. Gene ontogeny (GO) analysis further supported functional divergence: ThyAd I was enriched for pathways including “Mitochondrial organization” and “Positive regulation of cold-induced thermogenesis”, whereas ThyAd II was enriched for “Epithelial mesenchymal transition” (**Figure 2B**).

**Figure 2.**
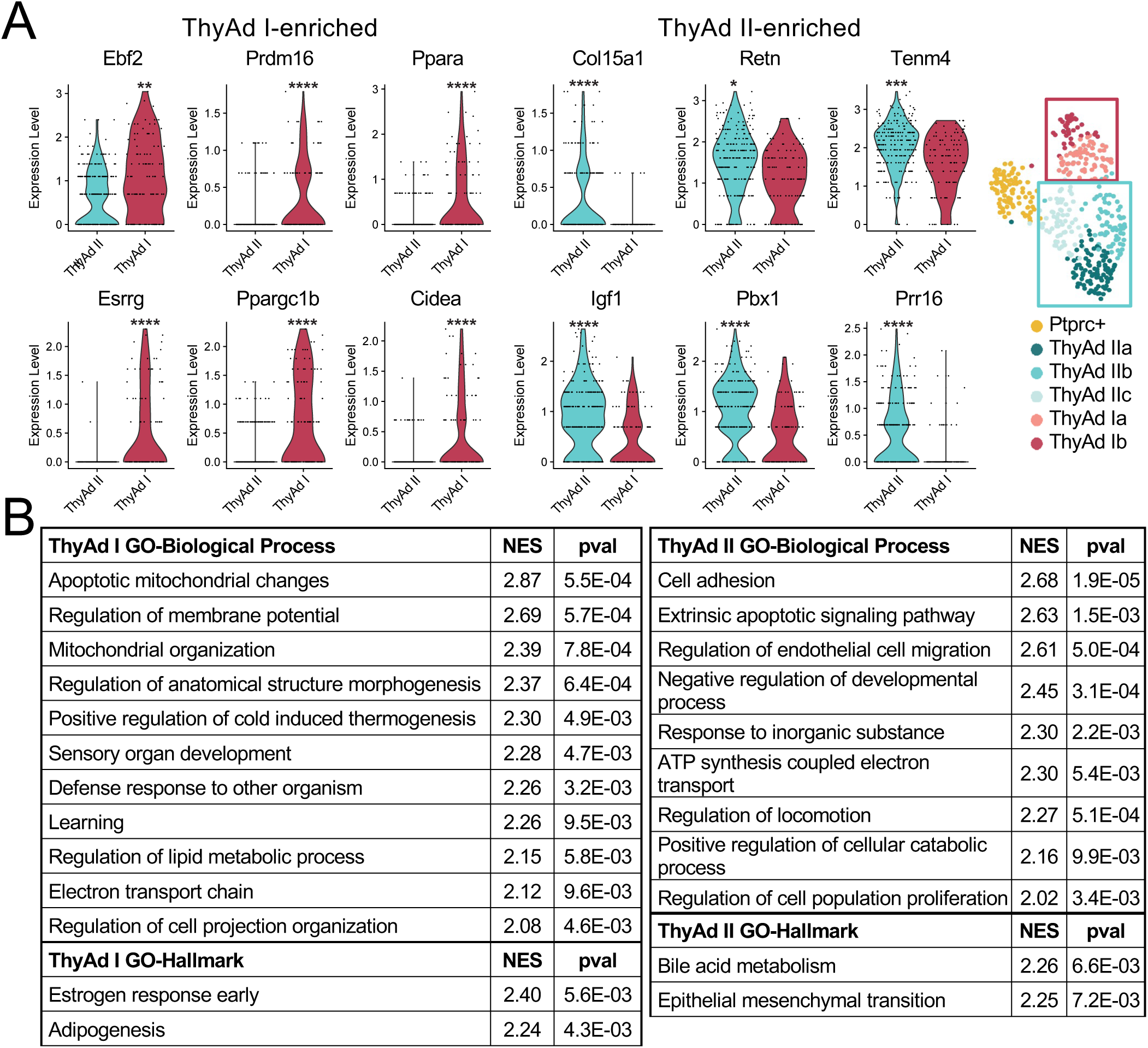
ThyAd I and II display highly differential gene expression. (A) Violin plots highlighting differential and shared expression of ThyAd I and II cluster-defining genes. Recolored UMAP denotes superclusters Ia/b and Ia/b/c are shown at right. (B) GO Biological Process Analysis for ThyAd Is and II superclusters.

### Single-nucleus ATAC-seq identifies state-specific chromatin accessibility and transcriptional factor motif enrichment in ThyAds

In addition to improving resolution of ThyAd states, the ATAC-seq component of the multiomic dataset enables inference of regulatory programs by identifying transcriptional factor (TF) motifs enriched in accessible chromatin. Moreover, because the chromatin state can precede or linger relative to active gene expression, the chromatin states can reflect transcription potential as well as recent enhancer decommissioning This can provide information on the possible antecedents for given populations because silent but recently decommissioned genes can retain areas of open chromatin in progeny cells. Because genomic DNA is fully preserved within the nuclei while much of the mRNA is lost, the ATAC-seq portion of the data provides high-quality information on gene expression states in isolated nuclei. We analyzed the ATAC-seq data focusing on genes associated with beige adipocytes: *Ppara*, *Cidea* and *Ucp1*. In all cases, increased chromatin accessibility was highly associated with the ThyAd Ib subcluster (**Figure 3A**). Additional genes are shown as expression and ATAC accessibility feature plots in **Figure S4**.

**Figure 3.**
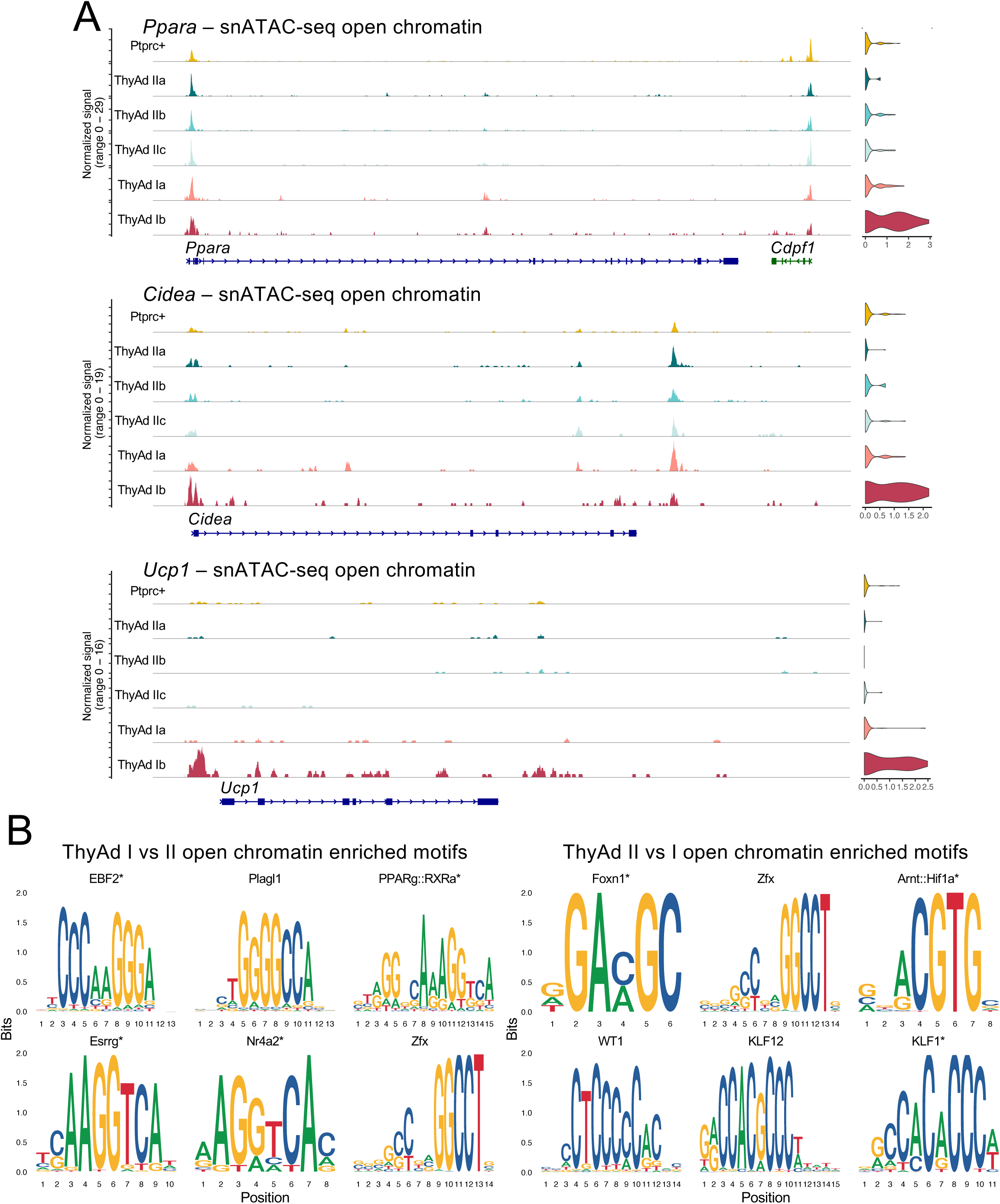
Differentially accessible chromatin and transcription factor motif enrichment in different ThyAd subpopulations. (A) Single-nucleus ATAC-seq open chromatin genome tracks at the *Ppara*, *Cidea* and *Ucp1* loci are shown for each subpopulation. (B) Motif analysis comparing combined ThyAd I and II subpopulations. The top 6 most enriched motifs are shown for each grouped population. Asterisks denote motifs unique to that population.

To identify candidate TFs underlying ThyAd state differences, we compared TF motifs enriched in between ThyAd I and ThyAd II. ThyAd I accessible regions were enriched for motifs consistent with adipocyte differentiation and beiging, including EBF, Plagl1, PPAR and Esrrg (**Figure 3B**). In contrast, ThyAd II accessible chromatin was enriched for Foxn1, hypoxia-inducible factor-1 (HIF-1), Wilms tumor protein 1 (WT1), Krüppel-like factor 12 (KLF12), and KLF1 motifs. Notably, Foxn1 is a master transcriptional regulator of thymic epithelial cells ^35^, suggesting a TEC-linked regulatory imprint in the ThyAd II population. This idea is consistent with lineage tracing experiments which identify Foxn1-expressing cells a progenitors of at least some adipocytes ^36^. Together with the EMT signature associated with ThyAd II gene expression (**Figure 1D and Figure 2B**), these results are consistent with a model in which ThyAd II white adipose cells arise from TECs. Consistent with this interpretation, transcripts encoding *Ebf2*, *Esrrg* and *Plagl1* were detected in subsets of ThyAd nuclei (**Figure S4A**). In contrast, other TFs highlighted by motif enticement (including *Foxn1*) were not detectably expressed yet displayed accessible regulatory regions (**Figure S4A-B**, **Tables S1 and S2**), consistent with residual chromatin accessibility following recent TF activity or enhancer decommissioning. Together, the data show increased accessibility at thermogenic loci and enrichment of EBF, PPAR, and ESRRG motifs in ThyAd I, while ThyAd II is enriched for FOXO1, HIF, and KLF1 motifs, consistent with a distinct and possibly TEC-linked regulatory imprint.

### Younger (4-month-old) male mice harbor ThyAd I and II populations distinct from gonadal fat

We also profiled ThyAds from 4-month-old male mice to align sampling with the earlier kinetics of thymic involution in males compared with females ^37–39^. Whereas 6-month-old females provided an optional window to capture ThyAd heterogeneity while thymic output remains relatively preserved, we reasoned that males would reach a comparable stage of remodeling at an earlier age. In mice, the thymus reaches near-maximal size around 5 weeks of age and then declines in cellularity. In male C57BL/6 mice, thymic cellularity decreases by approximately 50% by 4 months ^40^. This is matched in human thymic tissue 26-49 years old, where the perivascular space (PVS) comprises 50-60% of the thymic compartment and adipose tissue is easily identified ^2^. We therefore selected 4 months in males as a practical and biological window to capture ThyAds during an early-to-intermediate phase of involution.

We isolated ThyAd nuclei from 6 male mice maintained on standard chow and profiled them by single-nucleus RNAseq. From the same animals, we also collected gonadal (epididymal) white adipose tissue (gWAT) as a visceral fat comparator (**Figure 4A**). As in the female dataset, *Ptprc*+ nuclei carried through purification were manually removed prior to clustering. Joint clustering of ThyAd and gWAT nuclei revealed that the thymic and gonadal adipocyte transcriptomes were largely distinct, with ThyAd nuclei again segregating into two discrete clusters which neighbored but were distinct from specific gWAT populations (**Figure 4B**, gWAT II and III). Another gWAT cluster (**Figure 4B**, gWAT Ia/b). A rare thymic cluster expressed *Esrrg*, *Ppara* and *Ppargc1b*, consistent with a ThyAd I (beige-like) program (**Figure S5A**). In contrast, the ThyAd II cluster more closely aligned with one of the gWAT clusters, suggesting a more WAT-like transcriptional state (**Figure 4B** and **Figure S5A**). Feature plots confirmed strong expression of *Ppara* and *Esrrg* in ThyAd I (**Figure 4B-C**). These nuclei were also enriched for *Cidea* and *Ghr* (**Figure S5B**). When clustering thymic nuclei alone, ThyAd I and II remained clearly separable, marked for example by *Ppara* and *Ghr* (growth hormone receptor, **Figure 4D**). The most differentially expressed ThyAd I gene was *Ppara*, whereas ThyAd II was enriched for *Xdh*, *Fabp4*, *Adgrl4*, *Klf12* and *Tgfb2* (**Table S3**). Notably, *Tgfb2* (TGFβ2) has been implicated in promoting thymic involution ^41^. Although the different ThyAd clusters grouped more closely to gWAT clusters than to one another, we also assessed gene expression differences between the two tissues by grouping clusters and normalizing their gene expression. Among genes expressed in >5% of one or both nuclei with Adj. *p* < 0.05 and fold change >4 (529 genes), beige/brown adipocyte signatures were again a dominant program in ThyAds, as highlighted by enrichment of *Esrrg*, *Ppara*, *Ppargc1b* and *Cidea* in ThyAd nuclei (**Table S4**). Other genes globally enriched in ThyAds compared to gWAT included *Themis*, *Robo2*, *Dll4* and *Rasal2*. In contrast, gWAT-enriched genes included *Retn*, *Gpx1*, *Cfd*, *Lep* and *Apoe*. Both populations expressed core adipocyte genes including *Pparg* (**Figure 4D**) and *Cd36* (not shown).

**Figure 4.**
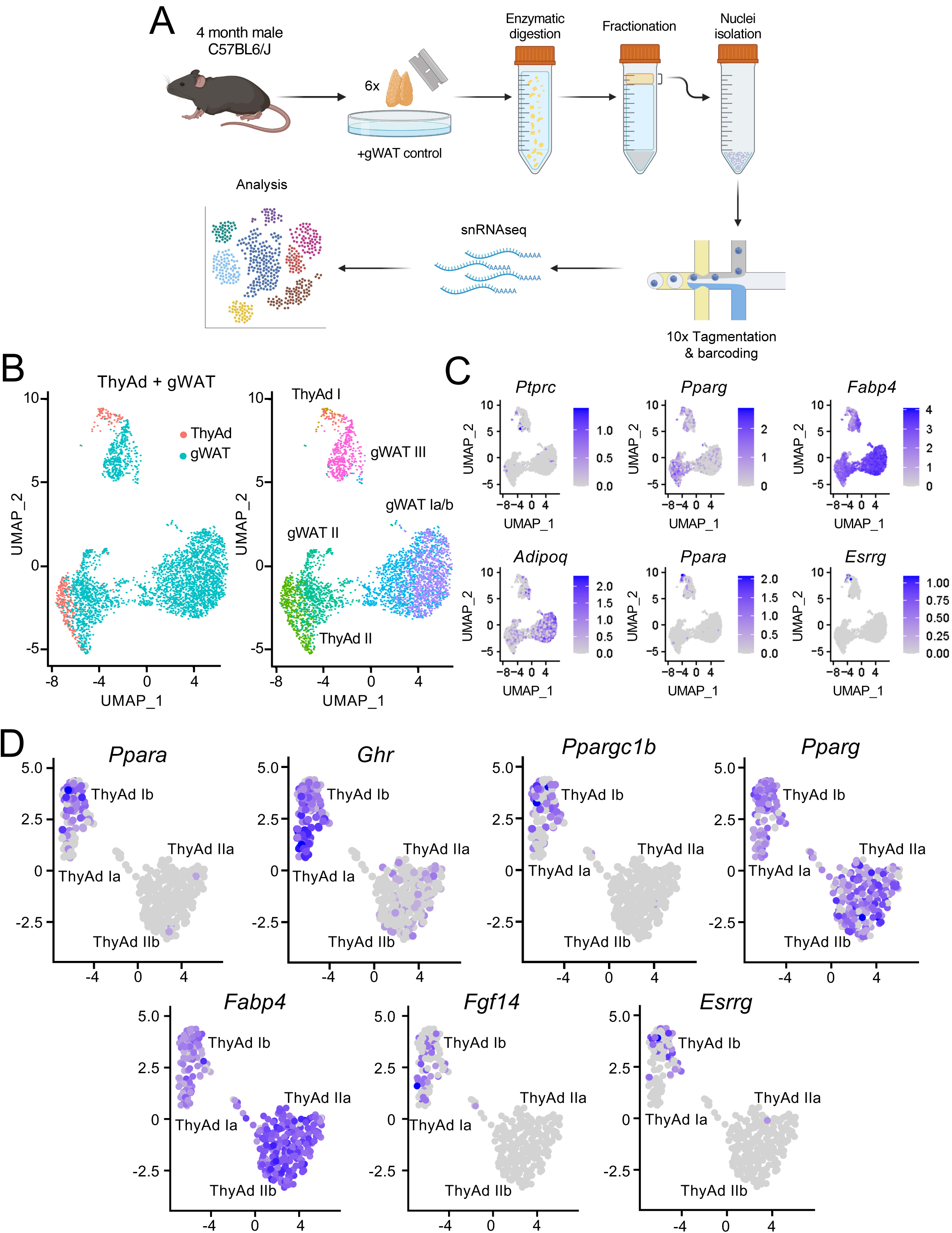
Male murine ThyAds are heterogenous and transcriptionally distinct from gWAT. (A) Experimental design for isolation and multiomic sequencing of murine ThyAds. (B) Single-nucleus RNA-seq UMAPs comparing ThyAds with gWAT isolated from 4-month-old male mice. Nuclei were labeled by tissue. (C) Feature plots showing expression of selected marker genes *Ptprc*, *Pparg*, *Fabp4*, *Adipoq*, *Ppara*, and *Esrrg*. (D) ThyAds were reclustered alone to distinguish 1a, Ib, IIa, IIb, and IIc populations. UMAP feature plots are shown for *Ppara*, *Ghr*, *Ppargc1b*, *Pparg*, *Fabp4*, *Fgf14*, and *Esrrg*.

### UCP1^+^ cells largely localize within the subcapsular cortical thymic compartment, whereas lipid-rich unilocular adipocytes localize to septa and perivascular space

Epicardial/perithymic fat is heterogenous, with some beige-like cells strongly expressing UCP1, a key enzyme that enables energetic equivalents to be consumed, indirectly generating heat ^42^. Although thymi used for sequencing were obtained from mice younger than 6 months old and were extensively cleaned of extrathymic fat prior to processing (**Figure 1B**), it remained formally possible that the beige-like ThyAd I signal reflected residual contamination outside the capsule. Prior work, however, has identified UCP1^+^ cells within the thymic parenchyma ^25^. To directly assess localization, we performed Immunofluorescence on 6-month-old female thymi. For this experiment, thymi were intentionally not cleaned of extrathymic fat, enabling direct comparison of perithymic adipose tissue with intrathymic UCP1^+^ cells.

As expected ^42^, perithymic fat and connective tissue abutting the thymic capsule was heterogenous, with regions containing large unilocular lipid droplets and others containing densely packed multilocular, BAT-like adipocytes with strong UCP1 staining. In contrast, UCP1^+^ cells within the thymic parenchyma were sparse and exhibited weaker, punctate UCP1. Omission of the primary antibody eliminated both the intra- and extrathymic UCP1 signal (**Figure 5**). In parallel, PPARγ antibody staining and BODIPY labeling highlighted adipocytes with large unilocular lipid droplets enriched within thymic septa and the capsule border, consistent with ThyAd II (**Figure S6**). Together, these data support the presence of beige-like ThyAd I cells within the thymic capsule and argue against perithymic contamination as the primary source of the ThyAd I transcriptional program detected by multiomic sequencing.

**Figure 5.**
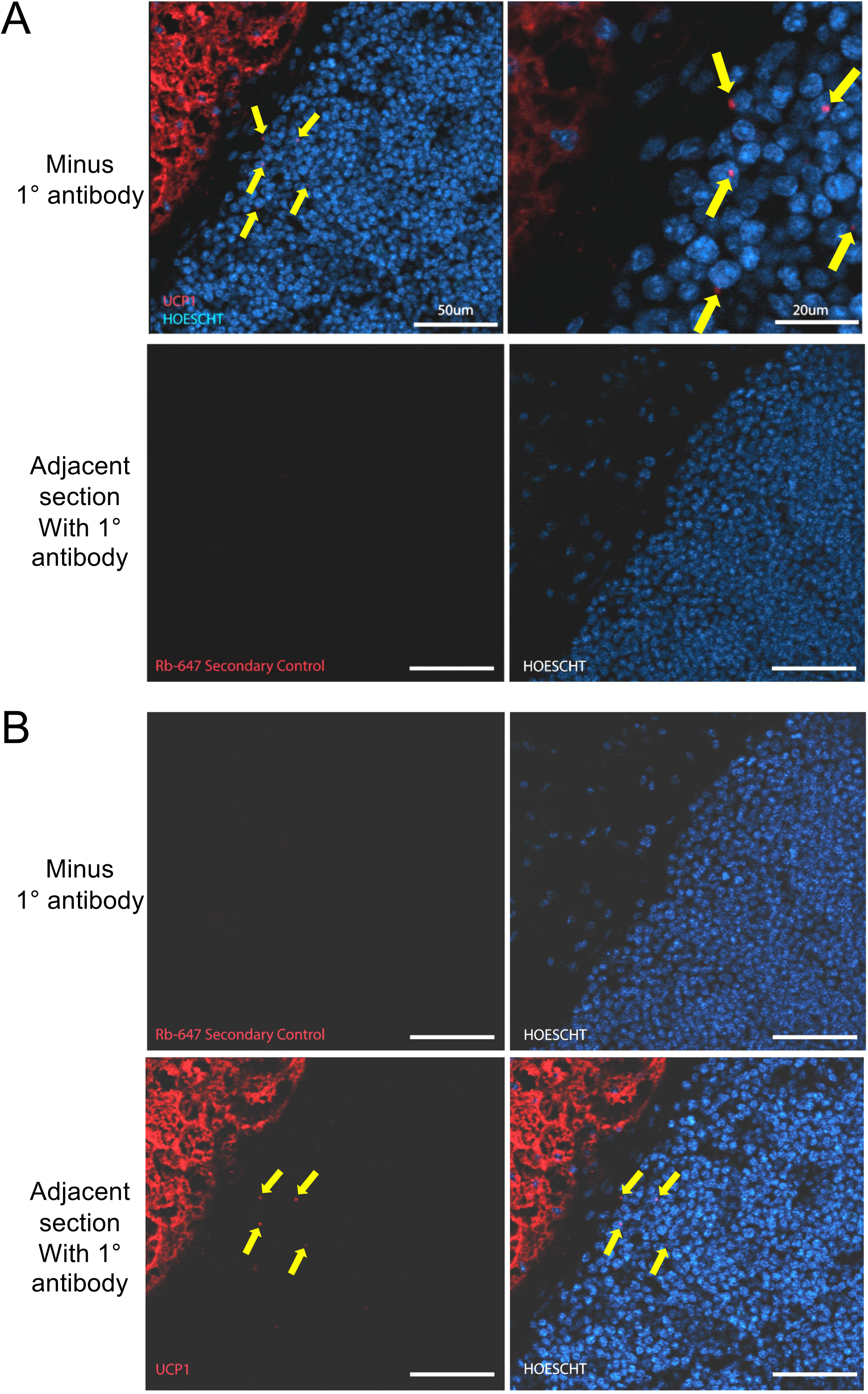
UCP1^+^ cells are detectable in the thymic cortical regions of 6-month-old female mice. (A) Top: 0.8 μm confocal Z-stack images of UCP1 immunofluorescence-stained cryosections of a region of a 6-month-old female murine thymus where the cortex meets the capsule. Slides were counterstained with Hoescht. Dense nuclear stain (blue) denotes the thymic parenchyma, predominantly TECs and thymocytes. Intense UCP1 staining is present in perithymic fat attached to the thymic capsule with a distinct expression pattern compared to scattered parenchymal UCP1+ cells (highlighted with yellow arrows). At right is shown a higher magnification image of the same slide. Bottom: an adjacent cryosection stained and imaged identically except lacking the primary antibody. (B) Similarly processed images of a different section from the same thymus.

## DISCUSSION

Recent advances in single-nuclei methodologies combining RNA-seq and ATAC-seq have been used to investigate the heterogeneity of adipose tissue depots throughout the body ^14,43^. Separately, they have been used to profile thymocytes and thymic stromal cells ^15–18^. However, adipocytes associated with the thymus (ThyAds) remain uncharacterized at a molecular level. Here, we developed methodologies to liberate and isolate adipocyte and their nuclei from mouse thymi, generating the first single-nuclei datasets of ThyAds. We identify similarities and differences in transcriptional profiles compared to a peripheral adipose depot. We profile nuclei from female and male thymi at two different ages, providing the first single-nucleus ThyAd atlas and expanding our understanding of ThyAds and their potential roles in the thymus and during thymic involution. By single-nucleus profiling, we identified two distinct ThyAd populations, termed ThyAd I and ThyAd II.

snRNAseq revealed that ThyAds express mature adipocyte-associated markers such as *Pparg*, *Fabp4*, and *Plin1*. No thymic preadipocytes were identified based on co-expression of markers such as *Pdgfra/b*, *Pref1* and *Cd34* ^44^. This is likely due to their relative lack of buoyancy. These studies also revealed a population of ThyAds expressing a highly metabolically active transcription program similar to beige adipose cells, termed ‘ThyAd Ib’. These nuclei were differentiated by significant expression of *Ucp1*, *Ppara* and *Esrrg*., hallmarks of beige and brown adipocytes. *Esrrg* for example encodes an orphan nuclear receptor expressed in metabolically active tissues (e.g., brain, heart, skeletal muscle, BAT) that promotes adipocyte beiging by activating mitochondrial biogenesis and oxidative metabolism ^27^. *Ppara* encodes a nuclear receptor associated with metabolically active tissues such as heart and brown adipose tissue (BAT) that oxidize fatty acids at a high rate ^45^. *Ppara* is also expressed in hepatocytes, where it promotes fatty acid oxidation and ketogenesis during fasting ^46,47^. In combination with PPARγ, PPARα can induce beiging of white adipose tissue ^48^. *Cidea*, which can be induced by the combined expression of PPARγ and PPARα and encodes a transcription coregulator that activates *Ucp1* expression ^28^. *Prdm16* and *Ucp1* expression are also hallmarks of BAT ^31^. PRDM16 represses type-1 interferon signaling in adipocytes which promotes thermogenic programming ^49^. Interestingly however, these nuclei did not strongly express other genes associated with beige adipocyte gene expression programs, e.g. *Tbx1*, *Shox2*, *Tmem26*, or *Hoxc9*, indicating that ThyAd Ib may be distinct from previously defined populations of beige adipocytes. An intermediate population – ‘ThyAd Ia’ – shared expression of beige/brown genes with ThyAd Ib such as *Ppara* and *Prdm16* but with little expression of *Ucp1*. DEG analysis of ThyAd clusters alone identified *Acsl1*, *Prkr2b* and *Mecom* among the topmost ThyAd Ia-enriched genes. The products of these genes are known to positively regulate adipogenesis and FA metabolism within BAT ^50–52^. ThyAd IIs displayed differential expression of genes such as *Col15a1*, *Prr16*, and *Pbx1*. These genes are often associated with EMT and ECM remodeling ^53–56^. EMT has been associated with more plastic and stem-like states, allowing de-differentiation and developmental plasticity, including the ability to transition to adipocytes ^57^. Within the thymus, age-associated TECs (aaTECs) are also associated with limited EMT signatures ^30^.

Analysis of transcription factor motifs enriched in open ThyAd chromatin supports the finding of two broad transcriptional ThyAd I and II programs and identifies potential regulators that may differentially regulate the two populations. Motif sequences for Esrrg, Ebf2 and Nr4a2 were highly enriched in ThyAd I, consistent with their significant expression in this population and roles in maintaining a beige-like transcription profile ^14,27,58,59^. Both Ebf2 and Esrrg have been demonstrated to promote and maintain a beige/brown transcriptional program ^27,60,61^. Esrrg accessibility was mostly restricted to ThyAd Ib in both male and female mice, suggesting Esrrg plays an important role in ThyAd Ib’s beige-like transcriptional state. Though *Ebf2* was expressed highly in all clusters, chromatin accessibility displayed a gradient pattern with higher accessibility in ThyAd Is and lower accessibility in ThyAd IIs. Interestingly, though Plagl1 motif accessibility was enriched in ThyAd Is, its expression by RNAseq was increased in ThyAd IIs.

ThyAd IIs instead showed increased accessibility for WT1, KLF12, and KLF1. WT1 is known to negatively regulate thermogenic gene expression and beige fat identity ^62^. WT1 also modulates EMT/MET during development ^63,64^. Interestingly, ThyAd II showed the strongest enrichment for Foxn1 motifs, which were not detected within the RNAseq. This is consistent with a model in which Foxn1 may have been expressed earlier in the etiology of ThyAd II. The origin of adipocytes within the thymic compartment is not well understood. Adipogenesis involves several tightly regulated steps, including the upregulation of PPARγ, a transcriptional factor required for activation of downstream genes ^65^. Aire-expressing TECs can give rise to “mimetic” cells that adopt characteristics of peripheral cells including antigens that would not otherwise be found in the thymus. These cells aid in thymic selection and the generation of central tolerance ^66^. In humans, Aire deficiency has been associated with Plin-1 autoreactivity and autoimmune adipose infiltration ^67,68^. Prior work has identified adipogenic potential in the thymic SVF ^36^. These findings suggest that at least a subset of the identified ThyAds may represent an uncharacterized depot of mimetic cells that eluded identification due to their buoyancy. Within the thymus, adipocytes have been suggested to arise from multiple sources, such as infiltration via proximity to the PVS or transdifferentiation from stromal cells with adipogenic potential ^69,70^. Lineage tracing experiments using LacZ reporters indicate that at least some adipocytes are derived from Foxn1-expressing TECs ^36^. We found that differentially open chromatin in WAT-like ThyAd II nuclei are enriched in binding sites for Foxn1, suggesting that this specific population may emerge from TECs. Other populations Foxn1-negative, EpCAM-negative stromal cells have been identified with adipogenic potential ^71^. Together with our findings of multiple distinct ThyAd populations, these findings are most consistent with a complex etiology for ThyAds.

We used confocal imaging to identify UCP1+ cells within the thymic parenchyma. These cells were sparse and well-spaced and mostly confined to cortical regions. These findings are consistent with work from Langhi et al., who identified several lipid-laden, Plin1+ populations including preadipocytes, brown adipocytes (UCP1+), macrophages (Iba1+), and pericytes (Ng2+) within the thymic parenchyma, surrounding blood vessels, and the subcapsular zone ^25^.

Thymocyte maturation is dependent on cellular crosstalk within the thymic microenvironment, including between thymic stromal cells and thymocytes. For example, TECs and fibroblasts secrete factors (e.g., IL-7, SCF, DLL4) that drive thymocyte proliferation/differentiation ^72^. Reciprocally, thymocytes provide TEC differentiation signals (e.g., RANKL, CD40, LTβ) to maintain thymic architecture ^1^. TEC-derived cells in the thymus are known to play additional roles beyond antigen presentation ^73^. More work is required to define ThyAd functions in antigen presentation and other functions such as providing trophic or metabolic support to thymocytes or TECs. Similarly, although ThyAds expand with age their role in thymic involution remains incompletely understood. Injection of T cell progenitors from young mice into aged thymi fails to rescue declining thymopoiesis. Conversely, transplantation of fetal thymic lobes into 18-month-old RAG1^-/-^ mice restores host thymocyte counts to that of 6-8 week-old mice ^74^. The fact that other strategies, such as injection of fetal TECs or ectopic expression of Foxn1 within adult TECs, only partially rescues aged thymi indicates that multiple attributes intrinsic to aged thymi are responsible for loss of T cell output ^75–77^. The possible contribution of one of more ThyAd subsets to this process also awaits experimentation.

### Limitations of the study

Our findings identify heterogeneity in ThyAds and identify beige-like cells expressing UCP1 within the thymic parenchyma. Although we confined the study to relatively young mice to reduce accumulation of epicardial/perithymic fat which could confound the study, and although thymi were extensively cleaned of fat outside the capsule before processing, it remains formally possible that one or more of the ThyAd subclusters could reflect contamination from perithymic fat, which is known to harbor beige-like cells. It is also possible that one or more subclusters could be molecularly indistinguishable from perithymic fat. Although prior lineage tracing work pinpoints TECs as a source for at least some ThyAds, and although our work is consistent with a TEC origin for ThyAd II, this has not been formally demonstrated. Finally, the study is intended to provide the first molecular ThyAd atlas, and as such contains no information of the functions for these cells in T cell development or thymic involution.

## METHOD DETAILS

### Laboratory mice

All mice were C57BL/6J background (#000664, Jackson Laboratories, Bar Harbor, Maine, USA). Mice were housed on a 12h dark/light cycle in groups (maximum five mice per cage) with ad libitum access to water and standard chow diet. All procedures were performed in accordance with the University of Utah Institutional Animal Care and Use Committee (Protocol #00001553).

### ThyAd isolation

Intact thymi were quickly removed and transferred to a petri dish on ice with ∼300uL PBS to keep tissue moist. Thymic capsules were cleaned of directly attached perithymic fat and connective tissues, rinsed a final time with PBS, and transferred to a 15mL conical tube. Digestion Buffer (1mM CaCl2, 250mM sucrose, 1.5mM MgCl2, 10mM KCl, 15mM HEPES, 0.1% BSA) with DNase I (8 μg/mL, D513, Sigma Aldrich) and Collagenase II (2 mg/mL, C1764, Sigma Aldrich) in DMEM-F12 10% FBS was added to each sample and thymi were digested for approximately 10-15 minutes at 37℃, horizontally on a shaker set at 300 rpm. Once most of the thymic tissue was digested, an equal volume of FBS was added to each sample to quench the reaction, mixed by gentle pipetting. Samples were fractionated by an initial centrifugation at 300 × *g* at 4℃. The topmost white layer containing mature, floating adipocytes was transferred to a clean 15 mL tube (Figure 1). Then, 10 mL of Wash Buffer (DMEM-F12, 15mM HEPES, 10% FBS) were added to each sample and mixed by gentle inversion and/or trituration with a micropipette to prevent bursting of the adipocytes. Samples were left on ice for 10 min, followed by a second round of centrifugation at 150 × *g* at 4℃. The white floating layer was transferred to a 5 mL conical tube and washed with 4 mL Wash Buffer. Samples were left on ice for 10 min, and spun at 150 × *g* at 4℃. The floating fraction was collected and transferred to a clean 1.5 mL eppendorf tube. Without adding more media, samples were centrifuged at 150 *x g* at 4℃. Residual buffer was carefully removed from the bottom of the tube with a 5 mL syringe and 21-gage needle. The remaining white layer of packed adipocytes was transferred to a clean, prechilled 1.5 mL eppendorf tube for nuclei isolation. For the data shown in Figure 4, 6 male thymi were collected, combined, and processed similarly. Gonadal (inguinal) fat (gWAT) was prepared from the same mice in parallel as a quality control.

### Nuclei and 10X library preparation

To isolate nuclei, samples were incubated with 100 μL 0.1x lysis buffer for 3-5 min on ice as previously described (10X Genomics, Demonstrated Protocol CG000366 Rev D). After washing and centrifugation following the outlined protocol, nuclei were resuspended in diluted 1x Nuclei Buffer (10X Genomics, PN 2000207). The concentration of nuclei was determined by AO/PI using a Luna-FL (Logos Biosystems, Inc., Gyeonggi-do, Korea). Approximately 5,000 nuclei were immediately loaded onto a 10X Next GEM Chip J (10X Genomics, PN 1000230). Single-nuclei libraries were prepared using Chromium Next GEM Single Cell Multiome Reagent Kit A reagents (10X Genomics, PN 1000284) and assessed for quality using an Agilent D1000 ScreenTape. Libraries were sequenced on an Illumina NovaSeq X instrument targeting 100 million paired end reads (GEX) and 300 million paired end reads (ATAC) of 150 bp for each sample.

### Data analysis

For female thymi, reads were aligned to the *mm10* reference genome (package 2020-A) to generate feature-barcode matrices using cellranger (version 2.0.2). Analysis was performed as described in the Seurat WNN and Signac 10x multiomic analysis vignettes ^78,79^. Briefly, doublets with high feature count and low-quality nuclei were removed by applying thresholding filters for number of ATAC transcripts (nCount_ATAC >500 & <70,000), unique genes (nFeature_RNA >200 & <2,500), mitochondrial mRNA content (mt.percent_RNA <20%), nuclesome signal (<2), and TSS enrichment (>1). 3,610 nuclei were recovered using this method for downstream analysis (S1B). Utilizing the Seurat SCTransform pipeline, dimensional reduction (PCA) was performed on the top 3,000 most variable genes. Equivalent normalization of the ATACseq data was performed by latent semantic indexing (LSI). A ‘weighted nearest neighbor’ (WNN) UMAP was generated using the FindMultiModalNeighbors function, taking both RNAseq and ATACseq datasets into account for subsequent UMAP and cluster visualization. Ptprc-enriched clusters were identified and excluded from downstream analyses while ThyAds were identified by enrichment of adipocyte markers Pparg and Plin1 but lack of residual immune cell markers such as CD8, CD4, and Ptprc (S1C, S1D). 429 remaining adipocytes were reclustered using the SCTransform pipeline.

For male thymi, 4,342 thymic and 4,968 gWAT nuclei were identified in each sample, respectively. Following removal of Ptprc-expressing cells, 1,521 filtered ThyAd nuclei and 3,806 filtered gWAT nuclei were present. Individual samples were normalized by SCTransform using the top 2,000 most variable genes followed by exclusion of immune cells as described above. Adipocyte clusters from individual samples were subjected to SCTransform and then merged by applying the Merge() and IntegrateLayers() functions. Merged dataset UMAPs were then generated as above by applying FindNeighbors() and FindClusters() with the standard pipeline. Subsequent filtering of adipocytes and removal of residual immune cells was performed as described above. After subsetting of adipocytes, SCTransform() was reapplied to normalize and scale expression of merged samples. Defining markers were again determined by applying the FindAllMarkers or FindMarkers function to identify differentially enriched genes in each cluster. Both datasets are hosted on the Gene Expression Omnibus (GEO) server under accession ID GSE319467. Cluster-defining markers were determined by applying the FindAllMarkers or FindMarkers function, which use a non-parametric Wilcoxon rank sum test with Bonferroni correction on all normalized features. For applicable figures, cluster-enriched markers are reported as levels of significance denoted by *p < 0.05, **p < 0.01, ***p < 0.001, and ****p < 0.0001. We also compared combined ThyAD clusters against all gWAT clusters to identify differentially expressed genes by applying FindMarkers(). Gene expression was normalized for comparison between tissues using SCTransform(). Filtered genes had an average log2FC > 2 or < -2, and Adj. *p ≤* 0.05.

### Motif, transcription factor, and GO analysis

Motif analysis was performed using chromVAR and additional packages as outlined in the Signac motif analysis and Seurat WNN vignettes ^79^. Clusters were compared against each other to identify the topmost differentially accessible motifs (p.val <0.05). To identify transcription factors (TFs) which may underlie ThyAd cluster transcriptional state, TFs with both high differential expression and their corresponding accessible motifs were determined using Seurat’s ‘presto’ package, which uses a Wilcox rank sum test to calculate p-values for both datasets and returning only TFs with p.val <0.01. GO analysis was performed on ThyAd I and ThyAd II cluster DEGs with adjusted p.val <0.05 using the ‘fgsea’ R package for Biological Process and Hallmark gene sets ^80^.

### Immunofluorescence and confocal microscopy

6-month old mice were euthanized by isofluorane and perfused with PBS. Intact thymi and directly attached connective tissue were carefully removed, washed in PBS, and postfixed for 30 min in 4% (w/v) paraformaldehyde (PFA). After 3x PBS washes for 20 min each, thymi were incubated in 10% sucrose for 1 hr and then left in 20% sucrose overnight at 4℃. The following day, thymi were incubated in a series of 30% sucrose for at least 2 hr, then incubated in 2:1 OCT compound:20% sucrose for 1 hr. Tissue was embedded in 100% OCT over a dry ice and 2-propanol slurry for sectioning. 10-20 μm cryosections were dried for 40-60 min at 37℃ and rehydrated in cold PBS. Tissue was rinsed 3x in PBST (0.1% Tween 20), and permeabilized in PBST for 15 min. Detergent was removed by rinsing 3x in PBS, following which samples were incubated in prewarmed 0.5-1X TrueBlack Lipofluscin Autofluorescence Quencher (Biotium, Fremont, CA) for 2 min. Slides were blocked for 1 hr at RT in PBS containing 10% normal donkey serum, 5% BSA, and 0.2% Triton-X, and incubated overnight in a primary antibody cocktail (10% donkey normal serum, 5% BSA) containing rabbit anti-UCP1 or rabbit anti-PPARγ (Cell Signaling Technologies, MA, USA) at 4℃. Sections were washed in PBS for 20 min thrice, followed by a 2 hr incubation with a secondary buffer cocktail at RT (1% BSA, 1:2000 Hoescht 33342, Thermo Fischer Scientific H3570) with or without donkey anti-Alexa Fluor 647 nm (1:400, Invitrogen, La Jolla, CA, USA). Slides were washed and mounted with Fluoromount-G (Thermo Fischer Scientific 00-4958). If substantial TrueBlack was removed from slides after incubation with blocking buffer, the post-staining treatment protocol was used per manufacturer’s protocol. Stained sections were imaged by resonance scanning confocal microscopy (A1R confocal system and Eclipse Ti inverted microscope; Nikon, Melville, NY, USA) with a 20X objective and 3X or 8X optical zoom (0.41µm/px).

## AUTHOR CONTRIBUTIONS

N.J.S. and D.T. conceived the study. KEV helped with generation of the ThyAd isolation protocol. N.J.S. and A.F. performed experiments. N.J.S. analyzed the data and generated figures. All authors contributed to the writing of the manuscript.

## ACKNOWLEDGEMENTS

We thank W.-L. Lo for critical reading of the manuscript. We thank M. Chandrasekharan for help with graphics. We thank B. Dalley, O. Allen and the University of Utah Huntsman Cancer Institute High-Throughput Genomics Core Facility. This work was supported by awards from the University of Utah Center Diabetes and Metabolism Research Center (DMRC), the NIH/NIAID (R01AI162929) and the Praespero Autoimmune Foundation (Canada) to D.T.

## SUPPLEMENTAL FIGURE LEGENDS

**Supplemental Figure 1. Quality control for single-nucleus ATAC-seq plus RNA-seq multiomics.**

(A) example nuclei following isolation. Areas of heterochromatin are preserved.

(B) Thresholding and quality control parameters employed to eliminate low-quality nuclei. Left: mitochondrial regression/normalization. Removed nuclei are shown in gray (top left). Right: Density plot showing retained nuclei by TSS enrichment and total peak fragments. nCount_ATAC quantiles are: 5% 6438.45; 10% 7834; 90% 29520.1. TSS. Enrichment quantiles are: 5% 5.47; 10% 5.8; 95% 8.96.

(C) WNN UMAP clustering of 3,610 nuclei isolated from the floating fraction of female 6-month old thymic tissue. Clusters 3 and 10 are highlighted (red circles) and identified as Ptprc^lo^ and Pparg^+^Plin1^+^. This subset was used for further downstream analysis.

(D) Selected marker genes used to identify immune populations and adipocytes are shown as violin plots by cluster.

**Supplemental Figure 2. Lipid-metabolism-related gene expression comparing whole sample and ThyAds.**

(A) Dot Plots of whole sample ThyAds showing expression of genes associated with lipid metabolism. Genes associated with lipogenesis (*Fasn*, *Acaca*, *Elovl6*, *Dgat1*, *Dgat2*, *Srebf1*, *Acly*, *Scd1*, *Scd1*) and lipolysis (*Lipe*, *Pnpla2*, *Mgll*, *Gk*, *Abhd5*) are shown.

(B) The same analysis was performed for subsetted Pparg+Fabp4+Plin1+ ThyAds and the residual Ptprc+ cluster.

**Supplemental Figure 3. Expression of selected markers by ThyAds and top DEGs by cluster.**

(A) Heatmap displaying the top 10 DEGs for cluster including the Ptprc+ nuclei.

(B) WNN UMAP projections of genes encoding selected adipokines (*Adipoq*, *Retn*, *Lep*, *Cfd*) and beige/brown adipocyte marker genes (*Cidea*, *Prdm16*, *Ucp1*, *Esrrg*, *Ppargc1a*, *Ppargc1b*). *Plin1*, *Fasn*, *Ebf2*, *Zfp423*, *Col15a1*, *Aldh1a1*, *Prr16* and *Rems3* are also shown.

**Supplemental Figure 4. Gene expression and open chromatin states of potential ThyAd transcription regulators.**

(A) Feature plots of WNN UMAPS showing expression and chromatin accessibility for additional genes. *Ebf2*, *Esrrg*, *Plagl1*, and *Foxn1* are shown. Gene expression plots for *Foxn1* could not be generated because Foxn1 was not expressed in any nucleus. The WNN plot from Figure 1C is recapitulated at right for comparison.

(B) Single-nucleus ATAC-seq open chromatin genome tracks at the *Foxn1* locus is shown for each subpopulation.

**Supplemental Figure 5. Gene expression differences associated with ThyAd and gWAT tissue from 4-month-old male mice.**

(A) Heatmap displaying the top 10 DEGs within each ThyAd and gWAT subpopulation is shown.

(B) Similar to Figure 4C except additional genes are shown: *Ebf2*, *Cidea*, *Prdm16*, *Ghr*, *Ebf1*, and *Aifm2*.

**Supplemental Figure 6. PPARγ-positive cells identified within the thymus.**

IF image of a 16 μm cryosection of a thymus from a 6-month-old female mouse. Samples was stained with BODIPY (488 nm), rabbit anti-PPARγ (647 nm) and Hoescht (405 nm). Dashed lines highlight demarcation between thymic parenchyma and capsule or septa.

