## Supplemental figures and text for "Molecular heterogeneity and differential metabolic signatures in thymic adipocytes"

Supplemental material includes:

Supplemental Figs. S1-S6

Supp. Fig. 1

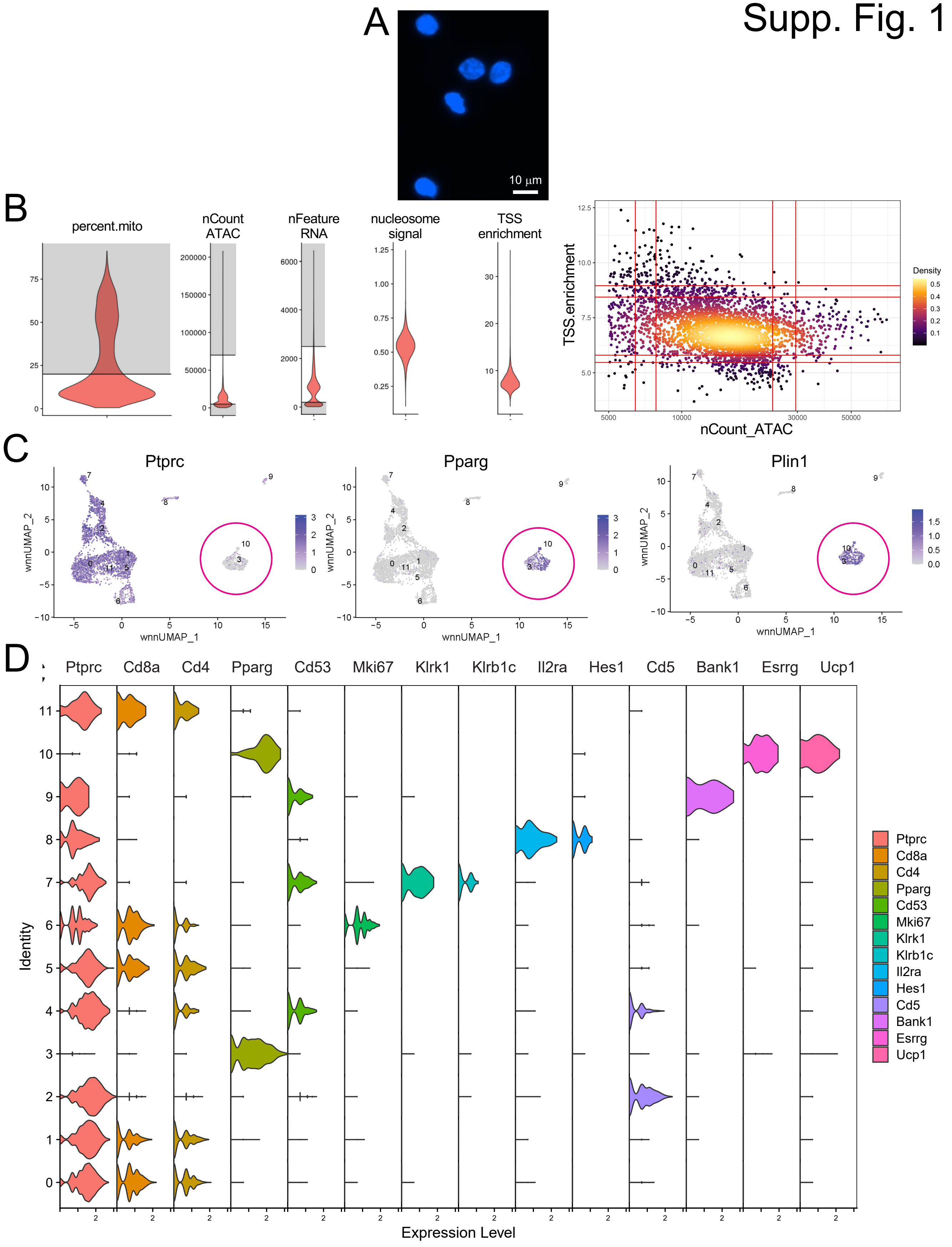
A
10 µm
B
percent.mito
nCount ATAC
nFeature RNA
nucleosome signal
TSS enrichment
TSS.enrichment
nCount_ATAC
Density
C
Ptprc
Pparg
Plin1
wnnUMAP_1
wnnUMAP_2
D
Ptprc
Cd8a
Cd4
Pparg
Cd53
Mki67
Klrk1
Klrb1c
Il2ra
Hes1
Cd5
Bank1
Esrrg
Ucp1
Identity
Expression Level

**Figure S1. Quality control for single-nucleus ATAC-seq plus RNA-seq multiomics.**

(A) example nuclei following isolation. Areas of heterochromatin are preserved.

(B) Thresholding and quality control parameters employed to eliminate low-quality nuclei. Left: mitochondrial regression/normalization. Removed nuclei are shown in gray (top left). Right: Density plot showing retained nuclei by TSS enrichment and total peak fragments. nCount_ATAC quantiles are: 5% 6438.45; 10% 7834; 90% 29520.1. TSS. Enrichment quantiles are: 5% 5.47; 10% 5.8; 95% 8.96.

(C) WNN UMAP clustering of 3,610 nuclei isolated from the floating fraction of female 6-month old thymic tissue. Clusters 3 and 10 are highlighted (red circles) and identified as $Ptprc^{lo}$ and $Pparg^{+}Plin1^{+}$. This subset was used for further downstream analysis.

(D) Selected marker genes used to identify immune populations and adipocytes are shown as violin plots by cluster.

Supp. Fig. 2

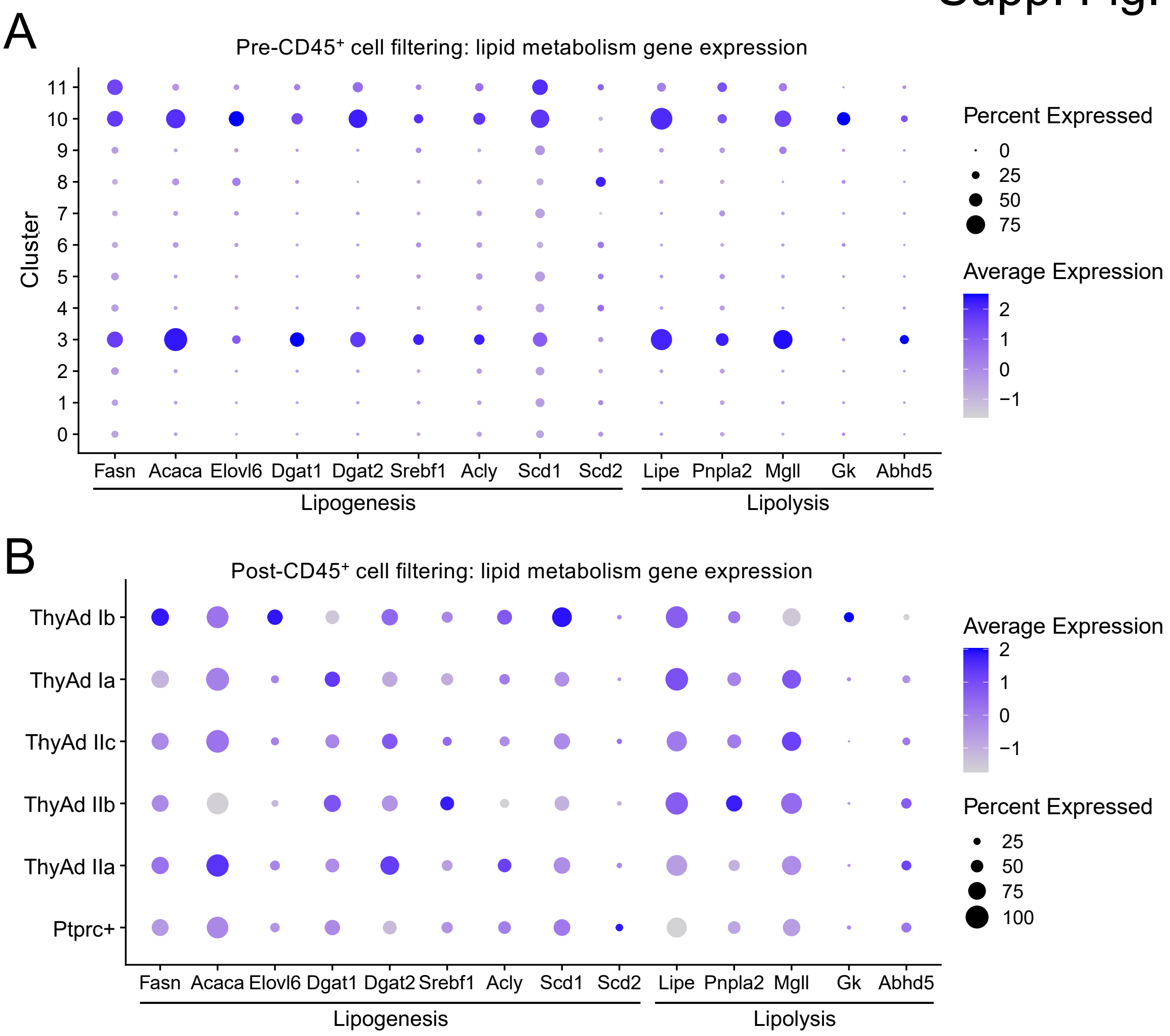


**Figure S2. Lipid-metabolism-related gene expression comparing whole sample and ThyAds.**

(A) Dot Plots of whole sample ThyAds showing expression of genes associated with lipid metabolism. Genes associated with lipogenesis (*Fasn*, *Acaca*, *Elovl6*, *Dgat1*, *Dgat2*, *Srebf1*, *Acly*, *Scd1*, *Scd1*) and lipolysis (*Lipe*, *Pnpla2*, *Mgll*, *Gk*, *Abhd5*) are shown.

(B) The same analysis was performed for subsetted Pparg+Fabp4+Plin1+ ThyAds and the residual Ptprc+ cluster.

Supp. Fig. 3

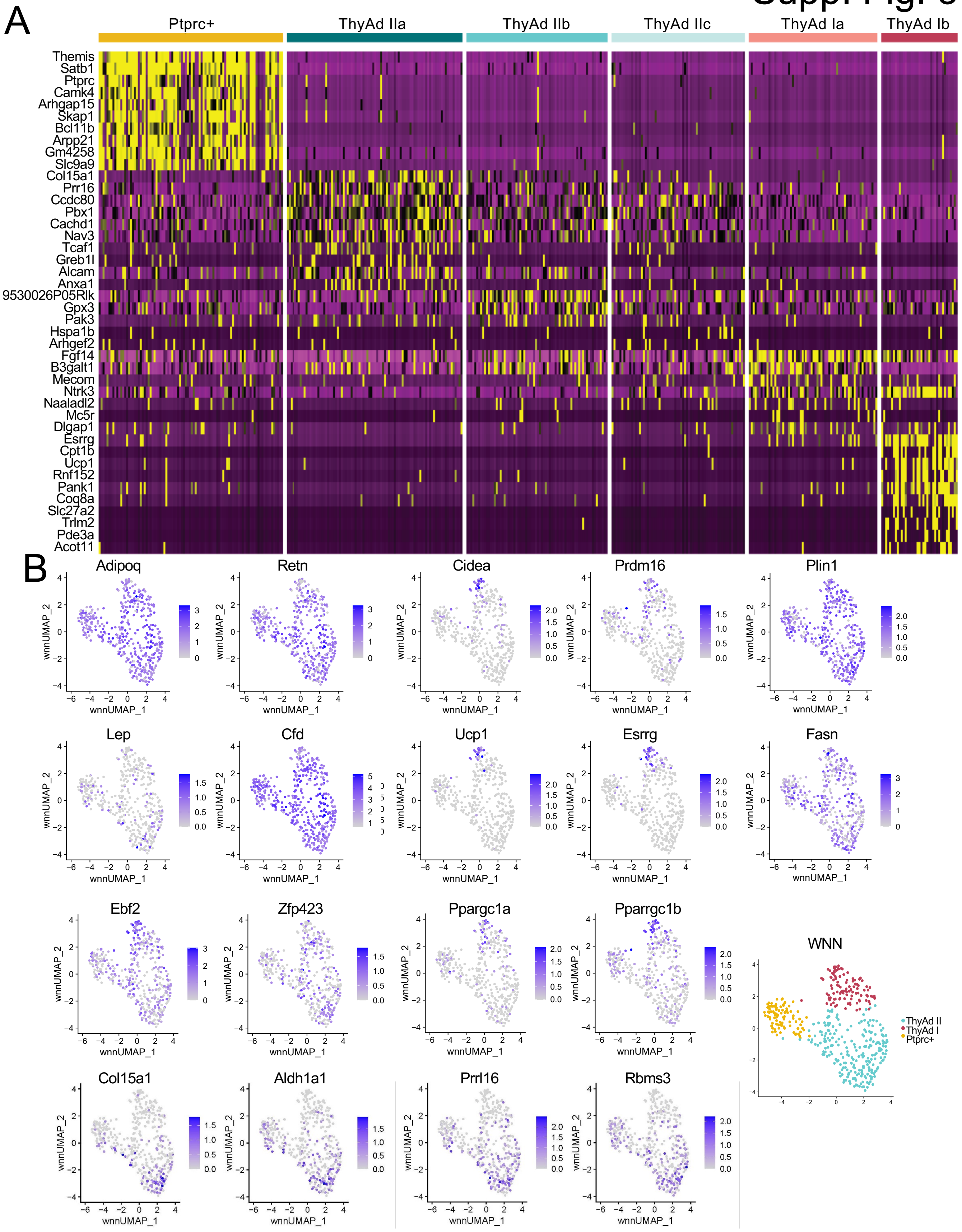
A
Ptprc+
ThyAd IIa
ThyAd IIb
ThyAd IIc
ThyAd Ia
ThyAd Ib
Themis
Satb1
Ptprc
Camk4
Arhgap15
Skap1
Bcl11b
Arpp21
Gm4258
Slc9a9
Col15a1
Prr16
Ccdc80
Pbx1
Cachd1
Nav3
Tcaf1
Greb1l
Alcam
Anxa1
9530026P05Rik
Gpx3
Pak3
Hspa1b
Arhgef2
Fgf14
B3galt1
Mecom
Ntrk3
Naaladl2
Mc5r
Dlgap1
Esrrg
Cpt1b
Ucp1
Rnf152
Pank1
Coq8a
Slc27a2
Trim2
Pde3a
Acot11
B
Adipoq
Retn
Cidea
Prdm16
Plin1
Lep
Cfd
Ucp1
Esrrg
Fasn
Ebf2
Zfp423
Ppargc1a
Pparrgc1b
WNN
Col15a1
Aldh1a1
Prrl16
Rbms3
wnnUMAP_1
wnnUMAP_2
ThyAd II
ThyAd I
Ptprc+

**Figure S3. OCA-B deletion in T cells decreases the $T_{CM}$- and naïve-like $CD4^+$ and $CD8^+$ T cell populations in MC38 TME.**

(A) Heatmap displaying the top 10 DEGs for cluster including the Ptprc+ nuclei.

(B) WNN UMAP projections of genes encoding selected adipokines (*Adipoq*, *Retn*, *Lep*, *Cfd*) and beige/brown adipocyte marker genes (*Cidea*, *Prdm16*, *Ucp1*, *Esrrg*, *Ppargc1a*, *Ppargc1b*). *Plin1*, *Fasn*, *Ebf2*, *Zfp423*, *Col15a1*, *Aldh1a1*, *Prr16* and *Rems3* are also shown.

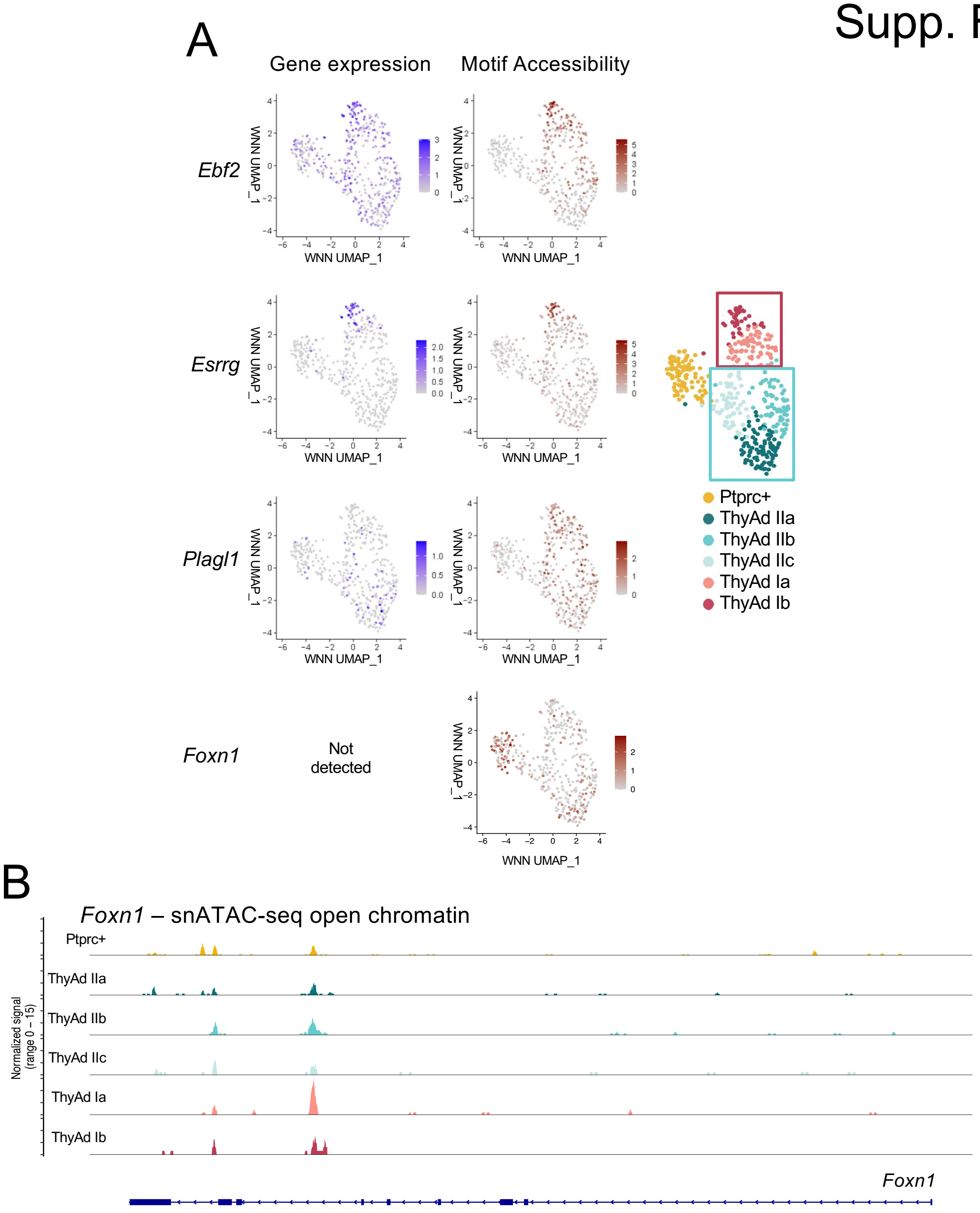


**Figure S4. Gene expression and open chromatin states of potential ThyAd transcription regulators.**

(A) Feature plots of WNN UMAPS showing expression and chromatin accessibility for additional genes. *Ebf2*, *Esrrg*, *Plagl1*, and *Foxn1* are shown. Gene expression plots for *Foxn1* could not be generated because Foxn1 was not expressed in any nucleus. The WNN plot from Figure 1C is recapitulated at right for comparison.

(B) Single-nucleus ATAC-seq open chromatin genome tracks at the *Foxn1* locus is shown for each subpopulation.

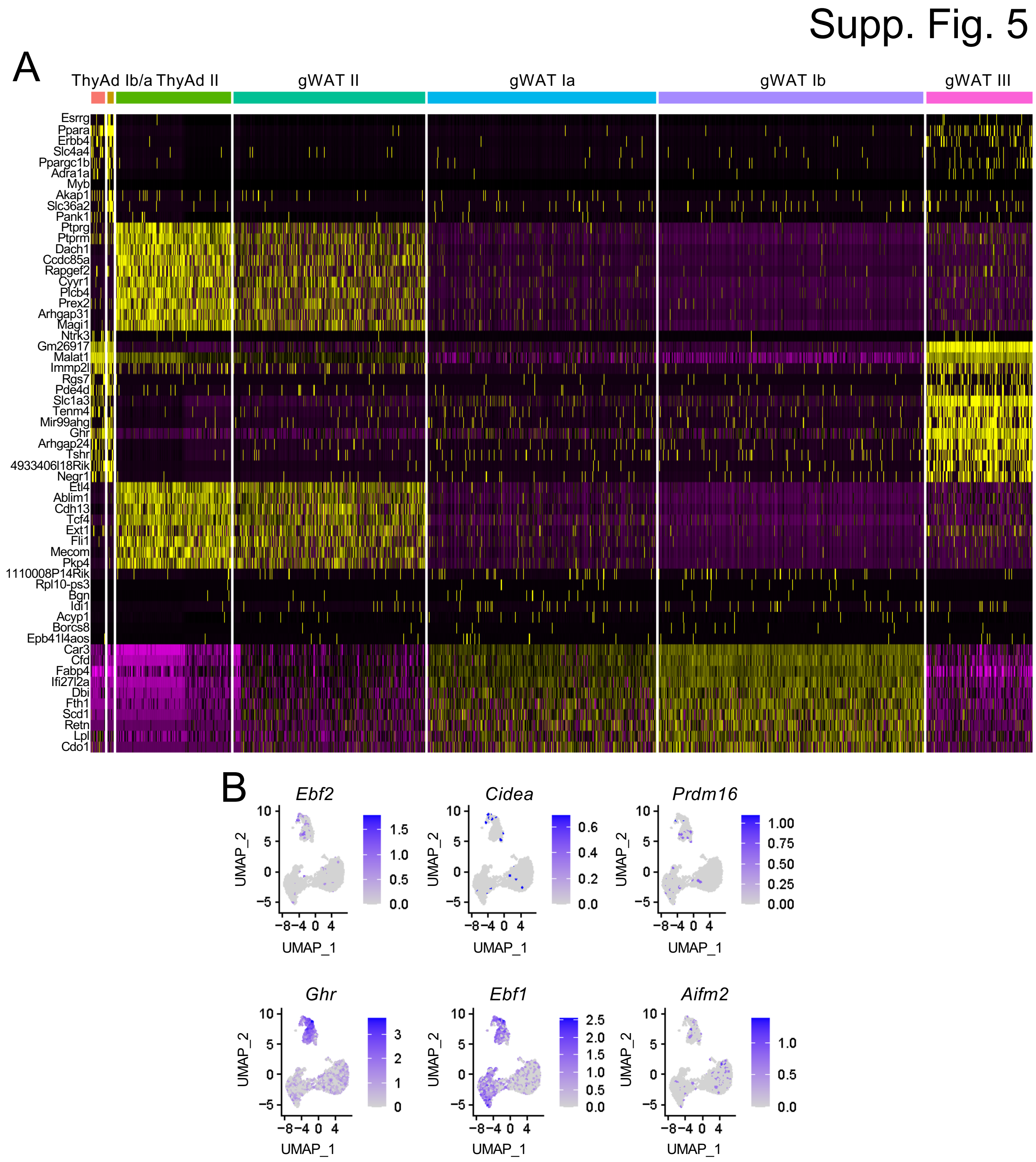


**Figure S5. Gene expression differences associated with ThyAd and gWAT tissue from 4-month-old male mice.**

(A) Heatmap displaying the top 10 DEGs within each ThyAd and gWAT subpopulation is shown.

(B) Similar to Figure 4C except additional genes are shown: *Ebf2*, *Cidea*, *Prdm16*, *Ghr*, *Ebf1*, and *Aifm2*.

Supp. Fig. 6

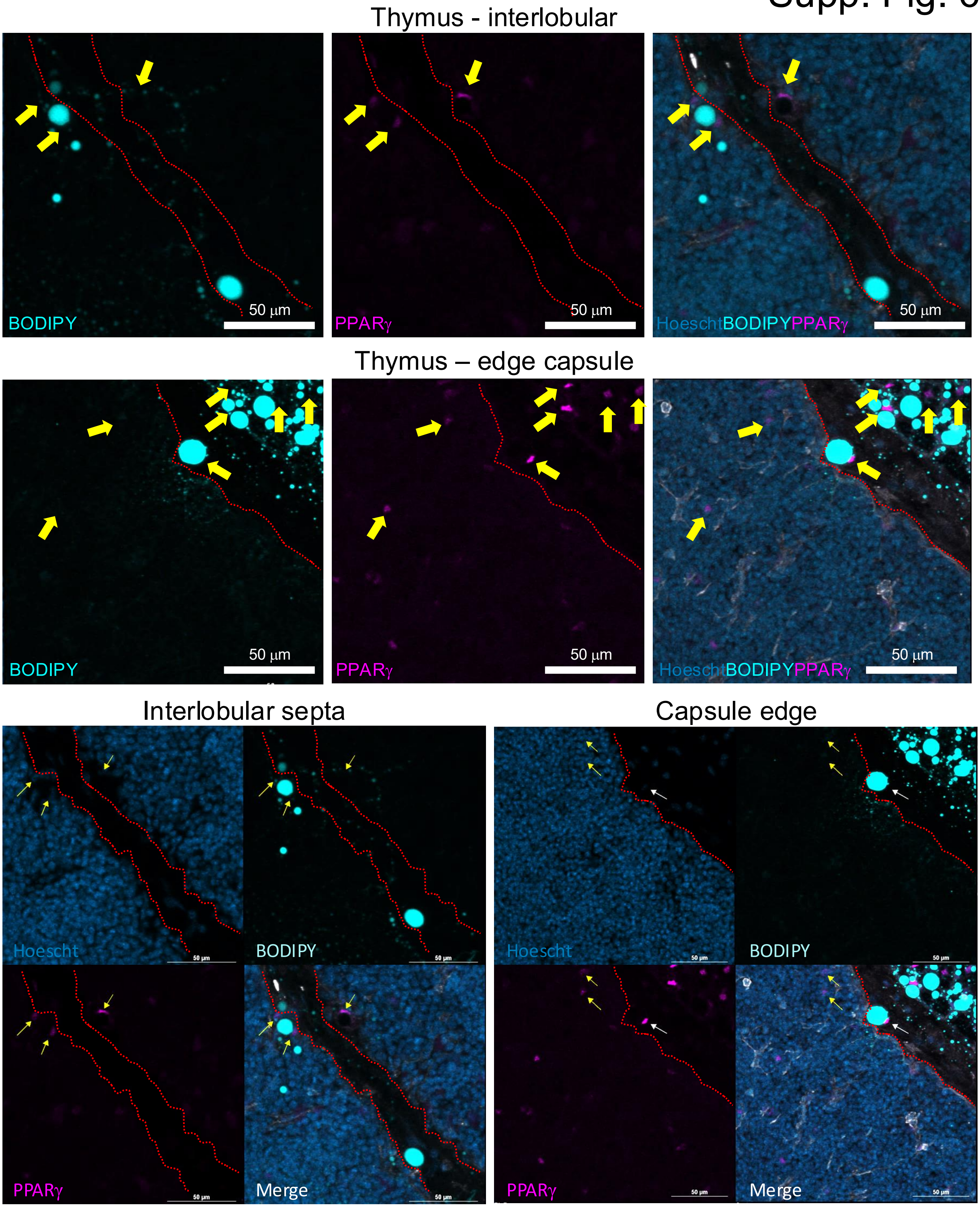
Thymus - interlobular
BODIPY
50 µm
PPARγ
50 µm
HoeschtBODIPYPPARγ
50 µm
Thymus – edge capsule
BODIPY
50 µm
PPARγ
50 µm
HoeschtBODIPYPPARγ
50 µm
Interlobular septa
Hoescht
BODIPY
PPARγ
Merge
Capsule edge
Hoescht
BODIPY
PPARγ
Merge

**Figure S6. PPARγ-positive cells identified within the thymus.**

IF image of a 16 μm cryosection of a thymus from a 6-month-old female mouse. Samples was stained with BODIPY (488 nm), rabbit anti-PPARγ (647 nm) and Hoescht (405 nm). Dashed lines highlight demarcation between thymic parenchyma and capsule or septa.
